# Differential effects of emotional valence on mnemonic performance with greater hippocampal maturity

**DOI:** 10.1101/2020.09.30.319178

**Authors:** Adam Kimbler, Dana McMakin, Nicholas J. Tustison, Aaron T. Mattfeld

## Abstract

The hippocampal formation (HF) facilitates the creation of declarative memories, with subfields providing unique contributions to the discriminability and generalizability of events. The HF itself and its connections with other structures exhibit a protracted development. Maturational differences across subfields facilitate a shift towards memory specificity, with peri-puberty sitting at the inflection point. Peri-puberty also happens to be a sensitive period in the development of anxiety disorders. Taken together, we believe HF development is critical to negative overgeneralization, a common feature of anxiety disorders. To investigate the role of the HF in behavioral discrimination and generalization we examined the relation between behavior and cross-sectional indices of HF maturity derived from subfield volume. Participants aged 9-14 years, recruited from clinical and community sources, performed a recognition task with emotionally valent (positive, negative) and neutral images. T1-weighted and diffusion-weighted structural scans were collected. Partial least squares correlations were used to derive a singular metric of maturity for both HF volume and structural connectivity. We found our volumetric HF maturity index was positively associated with discrimination for neutral images and generalization for negative images. Hippocampal-medial prefrontal cortex structural connectivity maturity metric evidenced a similar trend with behavior as the HF volumetric approach. These findings are important because they reflect a novel developmentally related balance between discrimination and generalization behavior supported by the hippocampus and its connections with other regions. Maturational shifts in this balance may contribute to negative overgeneralization, a common feature of anxiety disorders that escalates during the same developmental window.

**Significance Statement:** The hippocampal formation (HF) facilitates declarative memory specificity and is composed of subfields whose development during adolescence overlaps with the onset of anxiety disorders. Aberrations in mechanisms governing memory specificity may contribute to negative overgeneralization in anxious youth. Participants completed an emotional memory discrimination task while in the scanner. Using a multivariate maturity metric based on subfield volume we found individuals with more “mature” HF were better at differentiating similar neutral images and more likely to generalize similar negative images. These findings are important because they capture a novel developmental mechanism related to the balance between discrimination and generalization. Shifts in this balance, may contribute to negative overgeneralization, a common feature of anxiety disorders.

## Introduction

Memory increases in specificity, driven by maturation of key neurobiological substrate taking root around the onset of puberty and continuing into adolescence (Lavenex et al., 2007; Lee et al., 2014; Daugherty et al., 2017; Keresztes et al., 2018). Whether these specificity-supporting neurobiological mechanisms are similarly employed across different stimulus valences (e.g., emotional versus neutral) remains under-specified. Around the same developmental window, the prevalence of anxiety disorders increases (Beesdo et al., 2009). Together, understanding how developmental changes in memory specificity interact with emotional salience of stimuli may provide important insight into our understanding of negative overgeneralization, a characteristic symptom of anxiety where individuals generalize negative associations to similar events (Lissek et al., 2014). In the current study, we aim to understand the relation between cross-sectional indices of neurobiological maturation around the onset of adolescence and measures of behavioral discrimination and generalization to stimuli with emotional valence.

A network of interconnected regions in the medial temporal lobe (MTL) and medial prefrontal cortex (mPFC) govern the specificity of declarative memories. The MTL is comprised of the rhinal cortices, the amygdala, and the hippocampal formation (HF) (Squire et al., 2004). The HF can be further separated into distinct subfields: dentate gyrus (DG), Cornu Ammonis (CA) 1, 2, and 3, and subiculum. In human neuroimaging studies, the DG and CA3 subfields are often combined due to difficulties in reliably separating them (Daugherty et al., 2017), The amygdala is comprised of the central, medial, lateral, and basolateral complex nuclei (Phelps and LeDoux, 2005). Unique functional attributes are conferred by underlying architecture (Marr, 1971; Rolls, 2001) and connectivity (Ranganath et al., 2004; Goldstein et al., 2009; Bennett and Stark, 2016) of the MTL and mPFC. For example, the DG plays a disproportionate role in ‘pattern separation’ given its high number of granule cells and sparse firing, while the recurrent collaterals of the CA3 facilitate ‘pattern separation’ (Yassa and Stark, 2011; Rolls, 2013; Knierim and Neunuebel, 2016). The amygdala, contributes to the encoding of emotional salience of stimuli and events (Mcgaugh and Ayala, 2013) and the subsequent modulation of memory (McGaugh, 2004) with damage selectively impairing gist or generalized memories (Adolphs et al., 2001, 2005). While, the mPFC interacts with both the amygdala and HF regulating memory specificity (Colgin, 2011; Xu and Südhof, 2013; Jin and Maren, 2015; Sekeres et al., 2018).

Notably, the HF and mPFC are characterized by prolonged developmental trajectories extending well into adulthood with particularly rapid changes occurring around the onset of adolescence (Giedd et al., 1996; Lavenex and Banta Lavenex, 2013; DeMaster et al., 2014; Avino et al., 2018). Moreover, the distinct developmental trajectories have been shown to differentially impact behavior (Lee et al., 2014; Keresztes et al., 2018; Riggins et al., 2018). However, cognition does not emerge from the independent contributions of individual brain regions but rather reflects the product of networks of brain regions interacting (McIntosh, 2000). Thus, developmentally related increases in memory specificity likely reflect the composition of maturational changes across regions rather than the contribution of single regions alone(Keresztes et al., 2018).

To examine the role of MTL maturity on discrimination and generalization of stimuli with emotional valence in a sample of peri-pubertal youth, participants completed an emotional similarity task (Leal et al., 2014) where they rated the valence of stimuli (negative, neutral, positive) during a Study session in the scanner, then returned to the scanner after 12 hours for a surprise memory test where they made ‘old’ or ‘new’ judgments. Separate volumetric and connectivity based HF and amygdala maturity were assessed using a partial least squares correlation analysis (PLSC) using the structural and diffusion weighted scans (Keresztes et al., 2017, 2018). Behavioral measures of both discrimination and generalization were related to the multivariate measures of volume and connectivity. We tested the hypothesis that emotional valence of stimuli differentially produced increasing neutral discrimination and negative generalization with increasing hippocampal and amygdala maturity.

## Methods

### Participants

Fifty-two peri-pubertal youth (age 9-14 years) were recruited from mental-health clinics and the Miami-Dade community for a larger study to examine the neurobiological correlates of negative overgeneralization (McMakin et al., 2020). A strategy of recruiting from anxiety clinics and the community was used to maximize variability in key dimensions of interest—generalization and discrimination of emotional stimuli. Youth with anxiety are known to experience wide generalization gradients to negative stimuli in particular (Greenberg et al., 2013), which clinically appears as a tendency to pathologically extend fear from aversive contexts (e.g. house fire) to safe contexts with shared features (e.g. campfire). Participants recruited for the study were assessed for medical and psychiatric exclusionary criteria (e.g., current depressive episode, bipolar disorder, post-traumatic stress disorder, conduct disorder, oppositional defiant disorder, psychotic disorders, and obsessive-compulsive disorder) based on a screener assessing key symptoms associated with DSM-IV diagnoses and/or a parent-reported diagnosis. Following intake, three participants were ineligible for the study (left-handed) and another dropped out, leaving 48 volunteers to participate in the Study session scan. Two participants were not invited for the Test session scan due to excessive movement and an error in the experimental paradigm. A third participant failed to show-up for their appointment leaving 45 participants who completed the Test session scan. Eleven participants were excluded following the Test session: one failed to show up to their appointment, six for poor performance (hit rate for targets was 1.5SD below the average performance), three for errors in triggering the onset of the scanner with the task, and one participant was removed for excessive motion during the scan (defined as greater than 0.5mm of framewise displacement for more than 30% of volumes) leaving 34 participants (11.4 ± 2.0 years, 16 female) in the final sample. All participants provided written informed consent (legal guardian) and assent, and were compensated for their time.

### Behavioral Procedures and Methods

Participants took part in an emotional similarity task (Fig. 1). The task included an incidental encoding session during which participants viewed scenes (2000 ms) and were instructed to endorse images as either: negative, neutral, or positive. Stimuli were separated by a jittered inter-stimulus-interval (2000-6000 ms) during which a white central fixation was presented on a black background. Each scene was presented once, totaling 145 images (48, negative, 47 neutral, 50 positive). Participants returned 12 hours later for a surprise memory test, with approximately half (n=16) of them performing the task in the morning, post-sleep. During the Test session, participants were instructed to endorse images as either ‘old’ or ‘new.’ A total of 284 images were presented: forty-eight targets (16 each of negative, neutral, and positive) – repetitions of the images presented during the incidental encoding session; ninety-seven (32 negative, 32 neutral, and 33 positive) lures – images similar to but not exactly the same as an image shown during the incidental encoding session; and 139 foils (42 negative, 49 neutral, 48 positive) – images never presented before and not sharing similarity to the original images. Participants were asked to indicate whether each image was either ‘old’ (the subject recalls seeing that exact image during the Study session) or ‘new.’ They were instructed to endorse images as ‘old’ only if they were the exact same as the image seen during the Study session and to respond while the image was still on the screen. During the Test session each image was also presented for 2000 ms.

**Figure 1.**
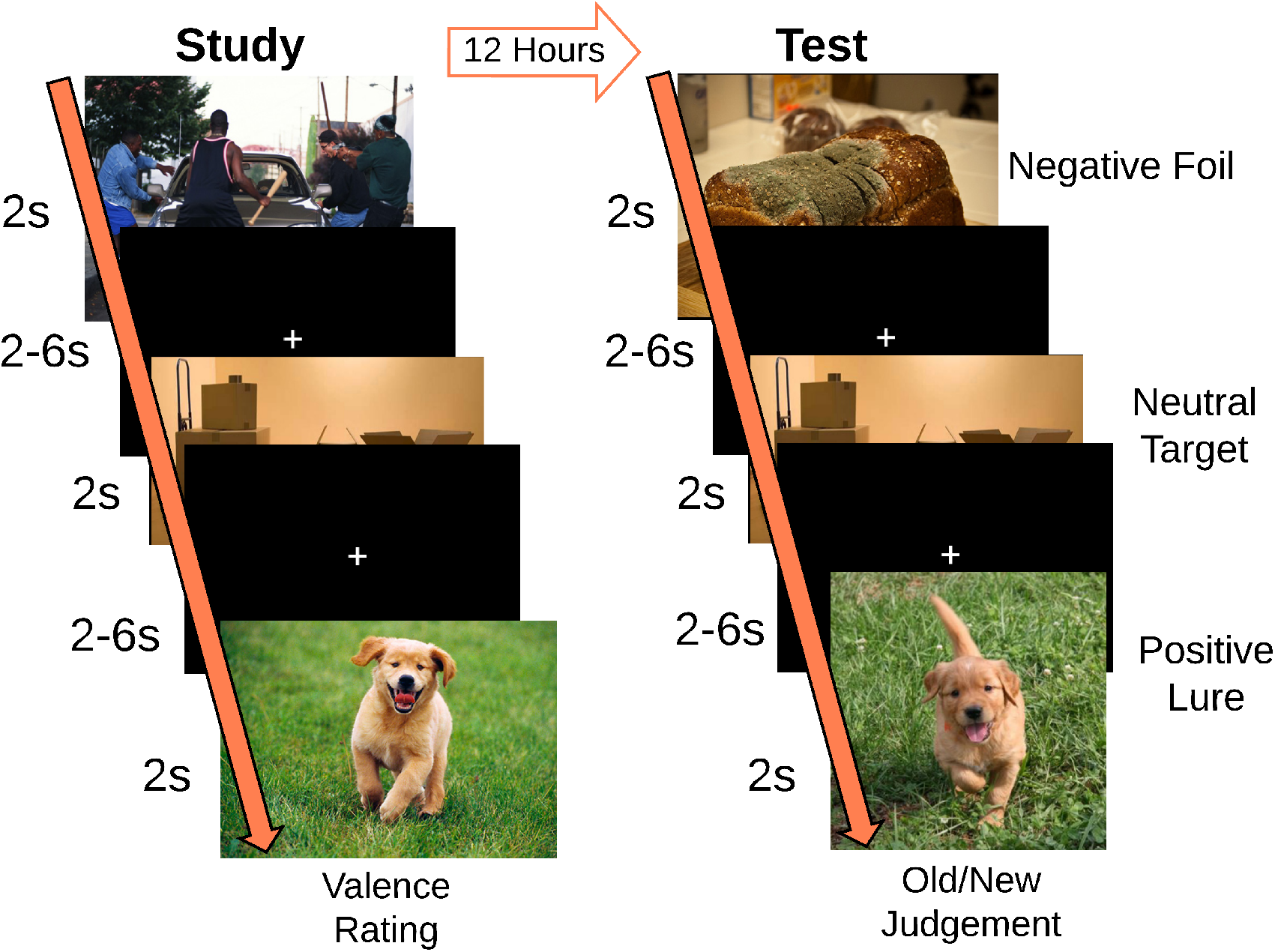
Trial structure of emotional similarity task. Participants experienced two scanning sessions. The Study session consisted of 145 images of differing valences while the Testing session consisted of a recognition memory task comprised of 284 images across the three valences with Targets (presented during the Study session), Foils (new images), and Lures (stimuli similar to the items presented during the Study session).

A lure generalization index (LGI) was calculated for each image valence by subtracting the proportion of old responses when given a foil image from the proportion of old responses when given a lure image (1).

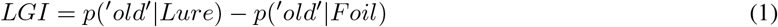

A lure discrimination index (LDI) was calculated for each valence by subtracting the proportion of new responses to targets from the proportion of new responses to lures (2).

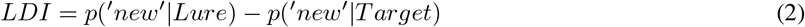

### Neuroimaging Data Collection and Preprocessing

Neuroimaging data were collected on a 3T Siemens MAGNETOM Prisma scanner with a 32-channel head coil at the Center for Imaging Science at Florida International University. Diffusion-weighted images (1.7mm isotropic) using a multiband sequence (slice acceleration factor=3, 96 directions, seven b=0 frames, and four b-values: 6 directions with 500s/mm2, 15 directions with 1000s/mm2, 15 directions with 2000s/mm2, and 60 directions with 3000s/mm2) in addition to a T1-weighted magnetization-prepared rapid gradient echo sequence (MPRAGE: TR=2500ms, TE=2.9ms, flip angle=8°, FOV=256mm, 176 sagittal slices, voxel size=1mm isotropic) were collected. A fieldmap opposite of the phase encode direction of the dMRI acquisition was also acquired for distortion correction.

FreeSurfer’s (version 6.0.0; Fischl, 2012) ‘recon-all’ algorithm was applied to each participant’s T1-weighted structural scan to obtain cortical surface reconstruction and cortical/subcortical segmentations. For diffusion image preprocessing, each participant’s diffusion weighted scan was registered to FreeSurfer structural space using boundary-based registration with the reference image being the first acquired b=0 frame. The FreeSurfer parcellation and segmentation file (aparc+aseg) was then binarized and transformed into diffusion space using FreeSurfer’s ‘ApplyVolTransform’ tool and was then binarized and dilated by 1mm to include edge voxels to act as a brain mask. Susceptibility distortion correction was then performed using FSL Topup, followed by eddy current correction using FSL Eddy on the diffusion data masked by the dMRI space brain mask. This preprocessed data was then input in to FSLs BEDPOSTX (Jbabdi et al., 2012) to model crossing fibers within each brain voxel. The results of BEDPOSTX were the basis of all subsequent probabilistic tractography based analyses.

### Delineating Amygdala Subregions

The amygdala is comprised of several nuclei, with each having unique anatomical connections to cortical and sub-cortical targets. We used probabilistic tractography combined with a novel method of k-means clustering analysis to identify amygdala subregions. Probabilistic tractography was computed from bilateral masks of the amygdala to 24 ipsilateral cortical and subcortical targets while avoiding the ventricles. This resulted in separate files (one for each target) containing the total number of random walks completed from each voxel in the amygdala mask to that specific targets. Each file was then vectorized and included in an n x m array, with n being the number of voxels in the left or right amygdala masks, and m being the number of amygdala targets. The array was then subjected to a k-means clustering algorithm with a limit of 4 clusters, with voxels serving as samples and targets serving as features, implemented in Python (scikit-learn). Each voxel in the amygdala masks were assigned a k-means cluster value based on their connectivity across the targets (aka, features), which were then coerced back into their three-dimensional anatomical representations. Volume estimates were then extracted for each of these clusters while correcting for ICV (Fig. 2).

**Figure 2.**
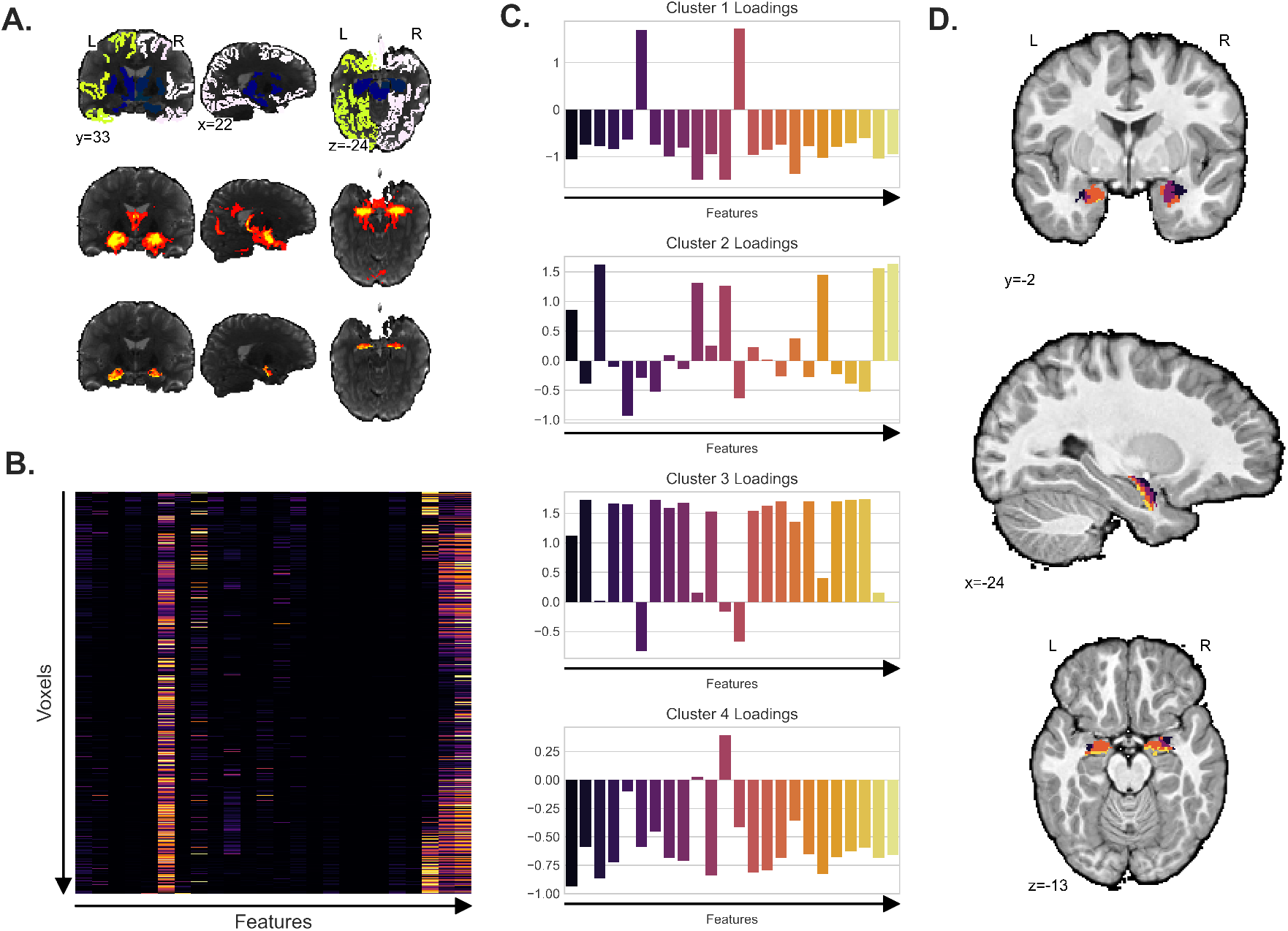
Delineation of the Amygdala through k-means clustering. (A) Probabilistic Tractography from the amygdala to 26 ipsilateral cortical and subcortical targets (first row) allowed for the definition of fiber pathways (second row) which allowed for the construction of metrics of anatomical connectivity in the form of number of walks to the target (third row; pictured is the number of successful walks from the amygdala to the HF). (B) From the random walk data, a matrix of number of random walks from amygdala to target was constructed across voxels in the amygdala. (C) The correlation matrix was then placed into a k-means clustering algorithm implemented in ‘scipy’ set to sort the data into 4 clusters, with cluster loadings pictured. (D). These four clusters were then classed for each participant, with an exemplar pictured.

### Delineating Hippocampal and Cortical Regions of interest (ROI)

Hippocampal subfields (bilateral DG/CA3, CA1, and subiculum) and the posterolateral and anteromedial entorhinal cortices were segmented using a consensus labeling approach. First, manual segmentations (Yassa and Stark, 2009; Yushkevich et al., 2015) were applied to an atlas set of 19 T1 MPRAGE scans and their corresponding T2-FSE scans (oblique orientation perpendicular to the long axis of the hippocampus; 0.47 mm2 in -plane, 2.0 mm slice thickness). Weighted consensus labeling from the atlas to an unlabeled T1 was accomplished by normalizing the atlas set to the unlabeled subject and applying multi-atlas segmentation with joint label fusion (Wang and Yushkevich, 2013). This approach capitalizes on both label and intensity information and has been used in a number of recent publications to segment hippocampal subfields (Sinha et al., 2018; Brown et al., 2019) (Fig. 3A). Cortical ROIs (e.g., perirhinal cortex, parahippocampal cortex, amygdala, superior frontal cortex, caudal and rostral anterior cingulate cortex, and medial orbitofrontal cortex) were created by binarizing FreeSurfer segmentations. All volume estimates were corrected for intercranial volume and age by multiplying each volume estimate by the ratio of age predicted whole-brain volume to actual whole-brain volume obtained via FreeSurfer.

**Figure 3.**
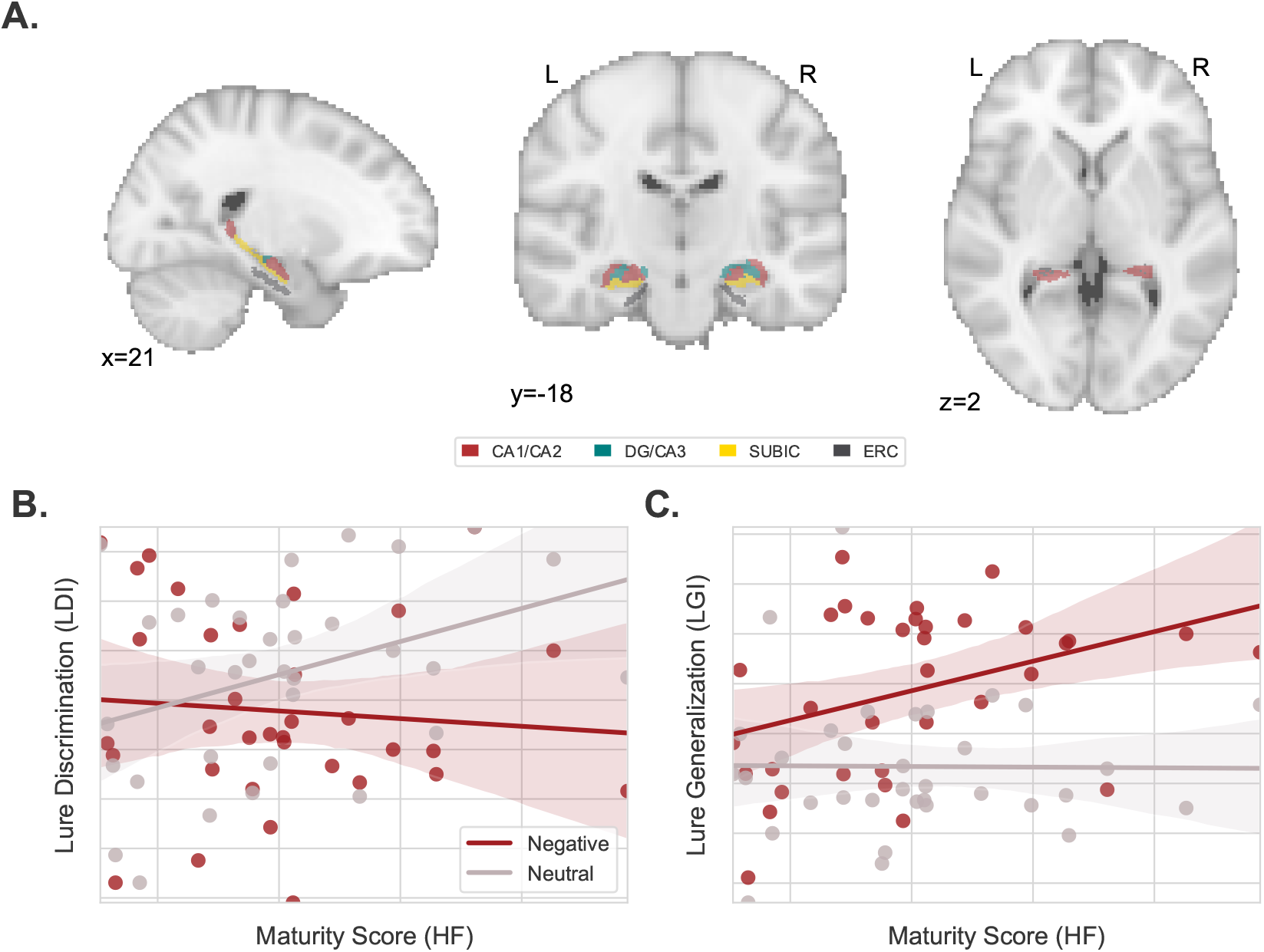
Differential relation between hippocampal maturity and memory performance across stimulus valence. (A) Hippocampal subfield volume estimates from the CA1/CA2, DG/CA3, Subiculum and entorhinal cortices (ERC) were used to produce a metric of hippocampal maturity as in Keresztes et al. (2017). (B) There was a significant interaction between image valence and hippocampal maturity predicting LDI performance (*maturity ∗ valence* : *F*(1, 32) = 6.159, *p* = .019, *R*^2^ = .161), with negative LDI showing no evidence of change across different levels of hippocampal maturity while neutral LDI performance increased. (C) An opposing significant interaction appeared for LGI performance (*maturity ∗ valence* : *F*(1, 32) = 5.737, *p* = .023, *R*^2^ = .126) where negative LGI increased across hippocampal maturity, whereas neutral LGI showed no evidence of change.

### Measures of Regional Connectivity

To examine the structural connectivity between regions, probabilistic tractography was conducted using FSL’s Prob-trackx (Behrens et al., 2003, 2007) with 25,000 streamline samples (step length=0.5, curvature threshold=0.2, maximum steps=2000) in each seed voxel to produce a connectivity distribution between each seed and target region, while avoiding paths through the ventricles. A list of the connections examined appears in Table 1.

**Table 1.** Seeds to target regions for connectivity analyses

| Connection | Seed Region | Target Regions |
| --- | --- | --- |
| Amygdala to HF | Amygdala | DG/CA3, CA1, ERC, Subiculum |
| HF to Rhinal | HF | PRC, pHPC, Posterior mERC, Anterior lERC |
| HF to mPFC | HF | Caudal ACC, Rostral ACC, Superior Frontal, mOFC |

### Regional Volumetric Maturity Estimates using PLSC

To calculate regional maturity scores partial least squares correlation (PLSC) was conducted to produce regional maturity for the HF (DG/CA3, CA1, subiculum, posterolateral and anteromedial entorhinal cortices), rhinal (RHI) cortices (posterolateral and anteromedial entorhinal cortices, perirhinal cortex, and parahippocampal cortex), and amygdala (AMY) (resulting k-means clusters determined by probabilistic tractography) according to an approach first outlined by Keresztes et al. (2017). A correlation matrix was produced by correlating AGE (in months) and volumetric measures for each ROI. The resultant matrix was then decomposed via singular value decomposition (SVD). Resultant weights for each ROI were then multiplied by each subject’s vector of ROI volumes to produce a singular regional maturity score. This process was completed for the HF, RHI and AMY to produce Hippocampal, Amygdala, and Rhinal Maturity Scores respectively.

### Connectivity Maturity Estimates using PLSC

PLSC was also used to compute connectivity maturity metrics between regions. A correlation matrix was constructed using AGE (in months) and the median number of random walks between the seed and target regions. This matrix was decomposed via SVD and the resultant weights were multiplied by the median connection strength to produce a connectivity maturity score. This was conducted for all connections outlined in Table 1. (Amygdala to HF, HF to Rhinal Cortices, and HF to mPFC).

### Statistical Analyses

Analyses into the effects of maturity and valence on lure generalization and discrimination outcomes were conducted using linear mixed-effects modeling conducted using the ‘lme4’ package in R version 3.6.1 (Bates et al., 2015). In this model valence and maturity were entered as predictors, and subject was modeled as the random intercept. Interactions were probed using simple-effects analysis also conducted in R.

## Results

### Hippocampal maturity was related to differential discrimination and generalization of neutral and negatively valenced scenes

To test our hypothesis that negative and neutral images experience different patterns of mnemonic discrimination and generalization with multivariate volumetric hippocampal maturity during the emotional similarity task we examined the association between individual’s HF maturity scores with their discrimination and generalization performance across image valence. Lure Discrimination and Lure Generalization Index (LDI and LGI respectively) scores were calculated (Equations 1 and 2) to assess a participant’s likelihood to discriminate or generalize lure stimuli, thought to be dependent neurobiologically on pattern separation and completion respectively. As multiple models are present in the current study, a Holm-Sidak corrected alpha of .026 was used to assess significance.

We observed differential patterns in the discrimination and generalization of emotionally valent lure stimuli across our HF maturity scores. As predicted, when comparing negative and neutral stimulus discrimination (e.g., LDI) we identified a significant interaction between HF maturity and stimulus valence (*maturity*∗*valence* : *F*(1, 32) = 6.159, *p* = .019, *R*^2^ = .161), with greater discrimination of neutral images associated with elevated HF maturity scores (*β* = .033, CI(95) = (0.003, 0.072), *t*(33) = 2.244, *p* = .032), while discrimination of negative images showed no evidence of such relationship with HF maturity (*β* = .016, CI(95) = (−0.041, 0.034), *t*(33) = −0.472, *p* = .640) (Fig. 3B). When assessing generalization (e.g. LGI) we similarly observed divergent patterns between negative and neutral stimuli (*maturity* ∗ *valence* : *F*(1, 32) = 5.737, *p* = .023, *R*^2^ = .126). We found enhanced generalization of negative images associated with greater HF maturity scores (*β* = .030, CI(95) = (0.003, 0.057), *t*(33) = 2.324, *p* = .027) while neutral generalization did not differ as a function of our HF maturity measure (*β* = −.000, CI(95) = (−0.023, 0.019), *t*(33) = −0.054, *p* = .957) (Fig. 3C)

### Valence related differences in discrimination and generalization not observed in the rhinal cortices or amygdala

To assess the specificity of the observed interactions with discrimination and generalization of emotionally valent images and our HF maturity scores, we probed for similar associations between maturity and emotional valence in neighboring MTL regions; such as the rhinal cortices and amygdala, two regions anatomically connected to the hippocampus (Suzuki and Amaral, 1994) and important for memory and emotional processing (McGaugh, 2004), with the rhinal cortices experiencing earlier development than both the amygdala and HF (Insausti et al., 2010). We did not observe divergent patterns in discrimination (LDI) across our multivariate maturity measures and image valence in the rhinal cortices (*maturity* ∗ *valence* : *F*(1, 32) = 0.000, *p* = 0.986) or the amygdala (*maturity* ∗ *valence* : *F*(1, 32) = 1.185, *p* = 0.284) (Fig. 4A,C); nor divergent generalization (LGI) across maturity measures and valence in the rhinal cortices (*maturity* ∗ *valence* : *F*(1, 32) = 0.690, *p* = 0.4123) and amygdala (*maturity* ∗ *valence* : *F*(1, 32) = 0.009, *p* = 0.925) (Fig. 4B,C).

**Figure 4.**
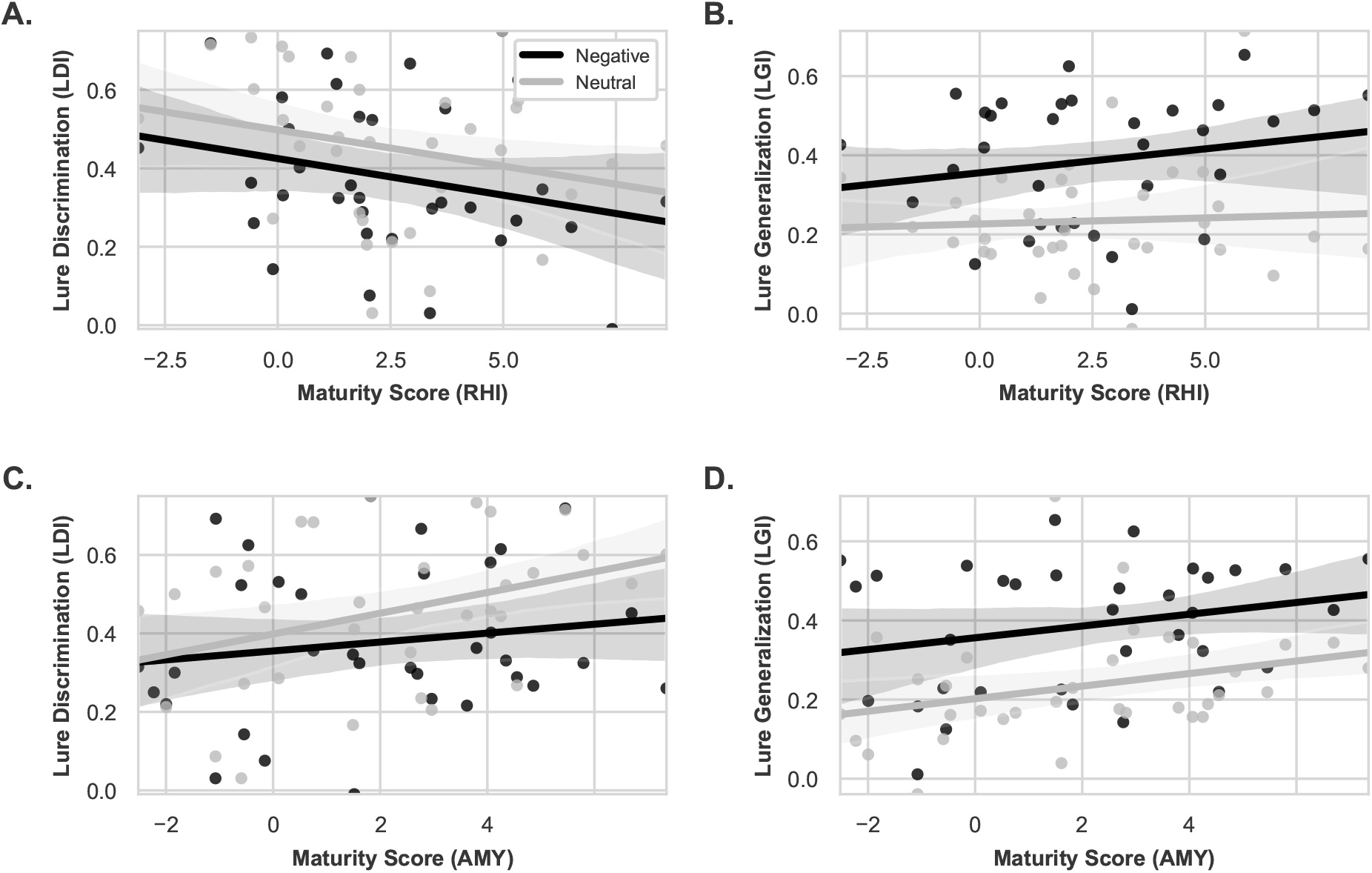
Rhinal cortices (RHI) and amygdala (AMY) show no differential relationship between maturity and valence with LDI and LGI performance. (A) The rhinal cortex maturity metric did not exhibit a differential relationship with discrimination for negative versus neutral valenced stimuli (*maturity ∗ valence* : *F*(1, 32) = 0.000, *p* = 0.986) (B) nor with generalization (*maturity ∗ valence* : *F*(1, 32) = 0.690, *p* = 0.4123). (C) The maturity metric for the amygdala similarly did not differentiate negative versus neutral stimuli for discrimination (*maturity* ∗ *valence* : *F*(1, 32) = 1.185, *p* = 0.284) (D) nor generalization (*maturity* ∗ *valence* : *F*(1, 32) = 0.009, *p* = 0.925) behavior.

### Hippocampal-mPFC anatomical connectivity exhibit trend towards differential relation between discrimination and generalization of emotionally valent stimuli

We next assessed whether non-volumetric measures of maturity, in this case diffusion weighted connectivity, were related to changes in discrimination and generalization with valence. We created a novel amygdala-hippocampal (AMY-HF) connectivity maturity metric (see Methods) and examined the associations between this measure and image valence with both LDI and LGI. We did not identify a divergence in discrimination (LDI) between negative and neutral images across our AMY-HF connectivity maturity measure (*maturity* ∗ *valence* : *F*(1, 32) = 1.243, *p* = 0.273) (Fig. 5A). When examining generalization (LGI) we similarly did not observe a divergence in generalization with image valence across changes in our AMY-HF connectivity maturity score (*maturity* ∗ *valence* : *F*(1, 32) = 0.034, *p* = 0.855) (Fig. 5B).

**Figure 5.**
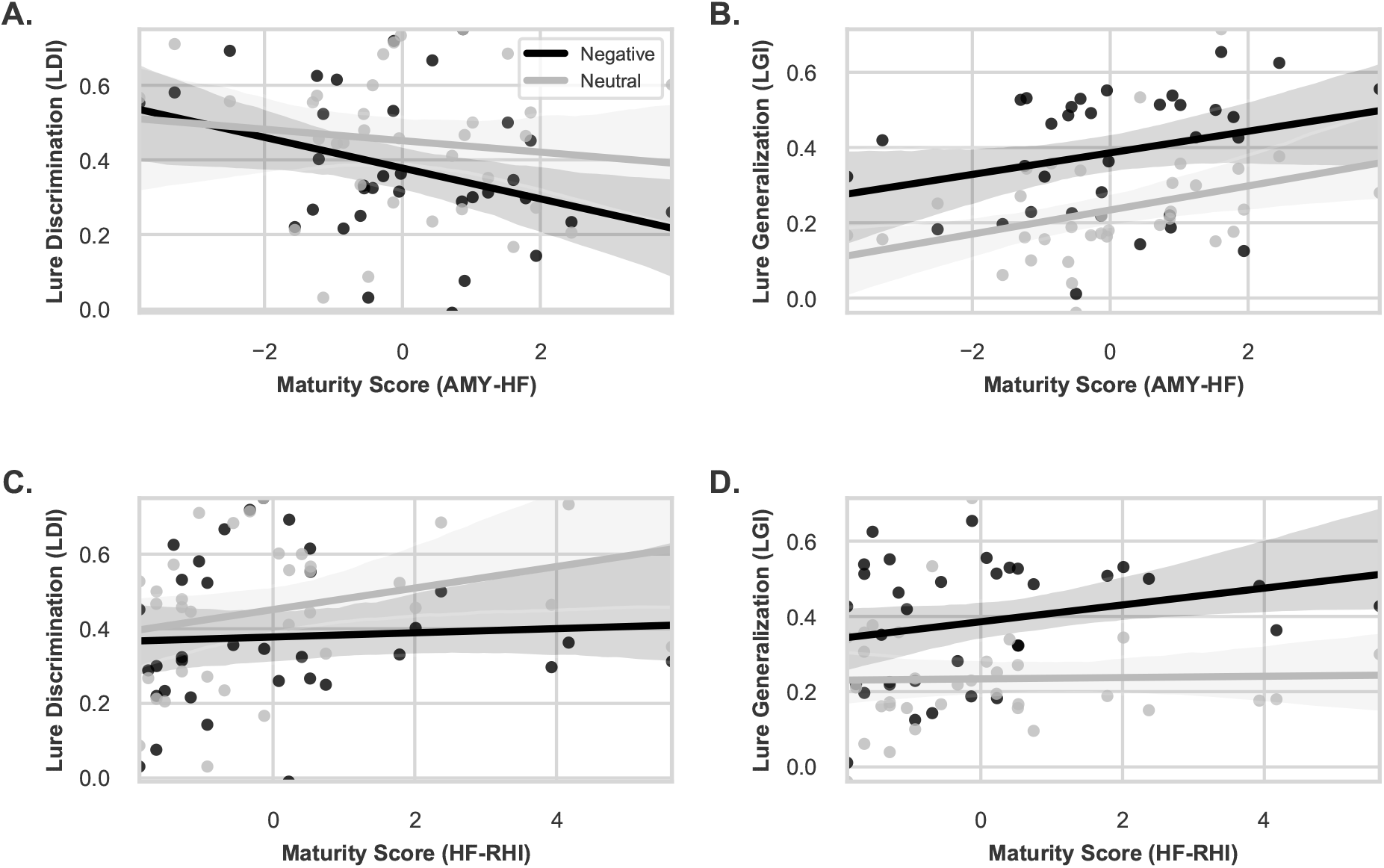
Association between connectivity maturity metrics and discrimination (LDI) and generalization (LGI). (A) AMY-HF connectivity maturity showed no differential relationship with LDI (*maturity ∗ valence* : *F*(1, 32) = 1.243, *p* = 0.273) or (B) LGI (*maturity ∗ valence* : *F*(1, 32) = 0.034, *p* = 0.855) across negative and neutral stimuli. Similarly, (C, D) HF-RHI connectivity maturity showed no valence related differential relationship with mnemonic performance (LDI: *maturity* ∗ *valence* : *F*(1, 32) = 1.312, *p* = 0.260; LGI: *maturity* ∗ *valence* : *F*(1, 32) = 1.833, *p* = 0.186).

We expanded this approach to other regions sharing anatomical connectivity with the hippocampus, creating maturity metrics for hippocampal-rhinal (HF-RHI) and hippocampal-mPFC (HF-mPFC) connectivity. We did not identify a difference between discrimination of emotionally valent stimuli across our HF-RHI connectivity maturity measure (*maturity* ∗ *valence* : *F*(1, 32) = 1.312, *p* = 0.260) (Fig. 5C). When examining generalization, we did not observe a divergence between emotional stimuli across our HF-RHI connectivity maturity measure (*maturity* ∗ *valence* : *F*(1, 32) = 1.833, *p* = 0.186) (Fig. 5D).

When examining the relationship between discrimination and generalization and our multivariate HF-mPFC anatomical connectivity maturity measure a familiar differential pattern was evident. When comparing discrimination across image valence (negative and neutral) and our HF-mPFC anatomical connectivity maturity score we observed a trend towards differential discrimination (*maturity* ∗ *valence* : *F*(1, 32) = 3.885, *p* = 0.057) (Fig. 6A). Generalization on the other hand exhibited an inverse trend (*maturity* ∗ *valence* : *F*(1, 32) = 3.095, *p* = 0.088) (Fig. 6B). These trending effects mirror the interactions observed in our volumetric HF maturity analyses (Fig. 3A,C).

**Figure 6.**
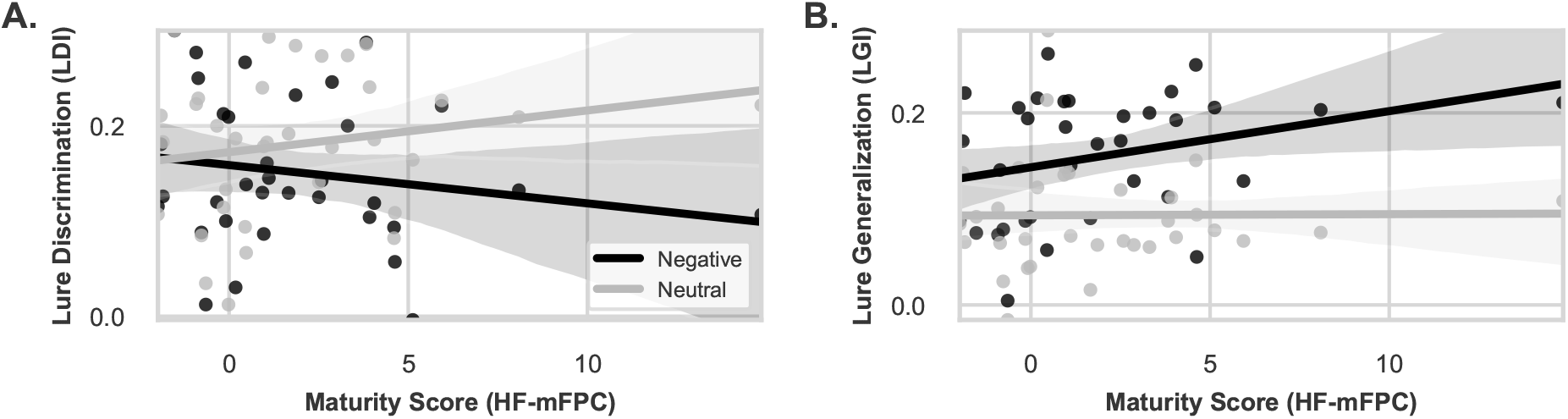
Maturity of HF-mPFC connectivity shows similar pattern of results to volumetric hippocampal maturity in relation to mnemonic performance. (A) There was a trending interaction between HF-mPFC connectivity and LDI (*maturity ∗ valence* : *F*(1, 32) = 3.885, *p* = 0.057), showing increased LDI for neutral and not negative images, (B) There was evidence of a trending effect of HF-mPFC connectivity predicting LGI (*maturity* ∗ *valence* : *F*(1, 32) = 3.095, *p* = 0.088).

## Discussion

We examined the role of MTL maturity (defined both volumetrically and through diffusion weighted connectivity) on discrimination and generalization of stimuli with emotional valence. We found greater volumetric maturity scores within the HF were related to enhanced mnemonic discrimination (LDI) performance for neutral images, as well as greater mnemonic generalization (LGI) for negative images. This effect of volumetric maturity appeared to be constrained to the HF and was not present in adjacent MTL regions (the amygdala and rhinal cortices). Employing the same multivariate decomposition approach, but using anatomical connectivity between the HF and other regions, a similar differential pattern of mnemonic generalization and discrimination across stimulus valence, while only trending, was evident in the connectivity between the HF and mPFC. Thus, development of the HF, assessed both volumetrically and through its connections to the mPFC, produce differential patterns of generalization and discrimination for negative compared to neutral images during a sensitive developmental period. This finding may have implications for our understanding of negative overgeneralization, a core feature of anxiety, which increases in prevalence during this developmental window (Beesdo et al., 2009; Lissek et al., 2014). Specifically, maturational changes in how emotional valence drives generalization may contribute to increasing negative overgeneralization and anxiety among vulnerable youth.

Previous studies have found positive associations between age related changes in the brain and mnemonic discrimination performance for neutral stimuli. For example, discrimination of neutral objects is positively associated with changes in DG/CA3 subfield volume across age (Lee et al., 2014). Similarly, greater volume of the CA2-4/DG was associated with enhanced memory performance for neutral trivia information in early childhood (Riggins et al., 2018). When using a multivariate decomposition derived from PLSC approach, a similar positive association between mnemonic discrimination of object images with HF maturity was shown (Keresztes et al., 2018). Our data corroborate these findings, showing increases in HF maturity, using PLSC derived maturity, are associated with improvements in mnemonic discrimination of neutral images. The current study also extends the findings from prior work, showing generalization of negative images is positively associated with HF maturity. Together, our results suggest that discrimination and generalization behavior are impacted differentially by emotional valence across age.

Generalization of emotional memories is adaptive and common. Behaviorally, learning under conditions of threat (Starita et al., 2019) or following negative (Schechtman et al., 2010) or aversive feedback (Resnik et al., 2011) facilitates generalization of episodic memories. While the amygdala has long been known to facilitate consolidation (McGaugh, 2004) and recently been shown to prioritize declarative memories through its coordinated neural activity with the hippocampus (Manns and Bass, 2016), damage to this region specifically impairs gist memories and leaves detailed memories spared (Adolphs et al., 2001, 2005). Neurons in the amygdala have been shown to have specific tuning properties related to generalization (Resnik and Paz, 2015) and specific populations in the lateral amygdala signal general versus cue-specific associations (Ghosh and Chattarji, 2015). While we did not identify in our results a difference in discrimination or generalization with emotional valence related to our volumetric or connectivity based metrics of maturation of the amygdala, a trend in amygdala volume (*F*(1, 32) = 3.679, *p* = 0.064) and connectivity with the HF (*F*(1, 32) = 5.373, *p* = .027) was associated with broad generalization irrespective of emotional valence. Rather, our results provide a novel contribution to the potential mechanisms underlying generalization and suggest the HF plays a unique developmental role, differentiating items to be discriminated from items garnering generalization. These results support the notion that the generalization of emotional stimuli is supported by a broad network of regions (Asok et al., 2019).

Differences in development across the HF contribute to heterogeneity of behavior well into adulthood. The HF experiences a protracted post-natal development in primates, with histology indicating profound increases in the DG volume (Lavenex et al., 2007). This protracted development has also been observed in studies of the human hippocampus using structural MRI. Some studies have reported greater hippocampal volume from childhood through adolescence (Ostby et al., 2009), while others have localized these changes to the hippocampal body and reported concomitant decreases of volume in the head of the hippocampus (Gogtay et al., 2006; DeMaster et al., 2014). Focus on individual subfields using structural MRI have identified increasing DG/CA3 volume as participants approach adolescence (Lee et al., 2014; Daugherty et al., 2017). Stereological studies in macaques have similarly demonstrated that DG granule cell populations mature well beyond early life (Jabès et al., 2011). Notably, the CA3 appears to mature in lock step with the DG (Jabès et al., 2011). Histological studies in humans have corroborated these findings, showing increases in postnatal volume across the HF marked by prolonged development of the DG and CA3, while the rhinal cortices exhibit a comparatively earlier maturational plateau (Insausti et al., 2010).

These differences in subfield volume and trajectories across development highlight the importance of using methods to simplify the input data when constructing models of HF development. This heterogeneity across the hippocampus is best captured by use of multivariate decomposition techniques (Keresztes et al., 2018). Our data demonstrates that findings such as those of Keresztes et al. (2018) are replicable across samples and different but related tasks using such techniques. Our data also demonstrates these techniques are sensitive to region specific changes, as indicated by different mnemonic outcome predictions between different constituent regions of the MTL (HF, amygdala, and rhinal cortices).

In addition to multiple internal structures, the HF shares robust anatomical connections with the mPFC (Varela et al., 2014). The mPFC contributes to schema development (van Kesteren et al., 2010) and is sensitive to information congruent with previous experience (van Kesteren et al., 2010). Developmentally related differences in the detection of congruence between previous and current contexts could influence generalization and discrimination behavior. The mPFC also plays an important role in mnemonic control, influencing memory specificity and generalization at encoding and retrieval (Xu and Südhof, 2013). When examining connectivity between the HF and mPFC using our PLSC metric, we found strikingly similar results to those in the initial analysis using only intra-HF volume.

The transition from childhood to adolescence (“peri-puberty”) is a neurodevelopmental window when neural networks associated with generalization (Bowman and Zeithamova, 2018) and emotional processing (Phelps and LeDoux, 2005) undergo dynamic change. At the same time, disorders of emotion, such as anxiety increase putting youth at a higher risk for escalating mental health problems (e.g. depression) in later adolescence and adulthood (Pine et al., 1998). Here we provide evidence for age related brain changes in the HF associated with the discrimination of neutral and generalization of negative stimuli. A developmental change in this balance may exacerbate negative overgeneralization in peri-puberty and ultimately help explain rising rates of anxiety in this developmental window.

Our study has several important limitations. First, our study was cross sectional, rather than longitudinal, and as such we can’t make any claims that these changes in volume with age are developmental in nature (Raz and Lindenberger, 2011). Second, HF subfields were defined using a consensus labelling approach. No harmonized protocol for HF segmentation exists, but are currently being devised (Olsen et al., 2019). Lastly, the limited sample size of the current study warrants caution of over interpretation and the need for replication in a larger sample.

Despite these limitations, our results both support and expand upon previous findings in the literature. Age related volumetric changes in the HF capture differences in generalization related to emotional salience and discrimination of emotionally neutral images. This difference in behavior appears unique to developmental changes in the hippocampus and may be related to changes in inter-regional connectivity between the HF and mPFC. Changes to the developmentally related balance between discrimination and generalization may support mechanisms of negative overgeneralization, a common feature of anxiety disorders often taking root during this developmental period.

## Acknowledgements

None

